# Arc capsid signaling between serotonergic and dopaminergic neurons sets sleep depth in *Drosophila*

**DOI:** 10.64898/2026.08.06.743357

**Authors:** Andrew R. Butts, Kylie DeNiro, Sven Bervoets, Jason D. Shepherd, Sophie Jeanne Cécile Caron

## Abstract

Arc genes evolved from Ty3 retrotransposons and encode proteins that self-assemble into virus-like capsids that package and transfer RNA between cells. This capsid-forming property may mediate a form of intercellular signaling distinct from classical synaptic transmission, but whether it operates within a defined neuronal system to control an ongoing behavior remains unclear. Here we show that Arc-dependent capsid signaling links serotonergic and PAM neurons to set sleep depth in *Drosophila melanogaster*. Loss of *dArc* genes deepened and consolidated sleep, increasing both sleep depth and the arousal threshold to mechanical stimulation, while leaving the diurnal sleep pattern intact. *dArc1* was required in serotonergic neurons and, to a lesser extent, in PAM neurons, and both the loss-of-function phenotype and its rescue depended on capsid formation. Knocking down *dArc1* in Sas-expressing cells, or Ptp10D in PAM neurons, reproduced the deeper, more consolidated sleep of *dArc^-/-^*flies, implicating the Sas-Ptp10D machinery in this signaling. These results identify Arc-dependent capsid signaling as a mechanism that links two modulatory neuronal populations to regulate an ongoing homeostatic behavior.

## INTRODUCTION

*Arc* genes evolved from the Ty3 family of retrotransposons and encode proteins that self-assemble into virus-like capsids^1,2,3,4,5^. These capsids package and transfer RNA, including their own mRNA, between cells^3,4,5,6,7^. This capsid-forming property may mediate a form of intercellular signaling in the nervous system that differs from classical synaptic transmission^5,8^. *Drosophila Arc* (*dArc*) genes, *dArc1* and *dArc2*, have been implicated in several behaviors and physiological processes: *dArc1* in the regulation of feeding and fat storage^9,10^, *dArc2* in long-term memory consolidation^11^, and *dArc1* in sugar reward valuation during learning^12^. It has been shown in non-neuronal tissues that Sas and Ptp10D, a ligand-receptor pair, deliver dArc1 capsids to their target cells^13^. For most of these functions, it is not known whether they require capsid formation or the Sas and Ptp10D machinery.

Studies established that *dArc2* is involved in memory consolidation^11^ and *dArc1* in sugar reward valuation^12^, and we showed that capsid formation is required specifically during sugar reward valuation^12^. *dArc1* is expressed in a subset of serotonergic neurons, and loss of *dArc* genes heightens the response of γ5 protocerebral anterior medial (PAM) dopaminergic neurons to sugar during reward valuation, supporting a model in which serotonergic neurons signal to PAM neurons through dArc1 capsids^12^. Whether capsid-mediated signaling is specific to sugar reward valuation or a general property of these populations of neurons is not known. Beyond reward valuation, which unfolds over seconds to minutes, both populations also regulate sleep, a state maintained over hours^14^. Serotonergic neurons promote sleep^15,16,17,18^, whereas PAM neurons promote arousal and oppose it^19,20,21^. It is possible that dArc1 capsids are used to mediate a signal between these populations beyond learning, in which case loss of *dArc* genes might affect sleep as well.

Here we show that Arc-dependent capsid signaling between serotonergic and PAM neurons sets sleep depth. Loss of dArc genes resulted in deeper and more consolidated sleep, while leaving the diurnal sleep pattern intact. *dArc1* was required in serotonergic neurons to set sleep depth, and this regulation depended on capsid formation. Knocking down *dArc1* in Sas-expressing cells, or *Ptp10D*, the Sas receptor, in PAM neurons, recapitulated the sleep phenotype of *dArc^-/-^* flies. Together, these results support a model in which dArc1 capsids transfer from serotonergic to PAM neurons, placing Arc-dependent capsid signaling within a defined population of neurons in the adult brain and showing that it regulates an ongoing, homeostatic behavior.

## RESULTS

### Loss of *dArc* genes increases and consolidates sleep but preserves the characteristic diurnal sleep pattern

To generate *dArc* mutants, we used CRISPR-mediated homologous recombination to delete either the *dArc1* or *dArc2* locus (*dArc1^-/-^* or *dArc2^-/-^*) (Figure S1A and B). We also analyzed a previously generated double mutant (*dArc^-/-^*) and a control line with an intact *dArc* locus (*dArc^+/+^*)^12^. We recorded locomotor activity in individual flies for four consecutive days using the multi-beam Drosophila Activity Monitor system. Sleep was inferred from periods of inactivity lasting five minutes or longer, as is standard in the field^22,23,24^.

*dArc^-/-^* flies slept substantially more than *dArc^+/+^* flies, at every hour of the day and night, increasing both daytime and nighttime sleep (Figure 1A and 1B). This increase reflected longer but fewer sleep episodes (Figure 1C and 1D), indicating that sleep in *dArc^-/-^* flies was more consolidated, while the normal diurnal sleep pattern — a relatively shorter daytime siesta followed by longer nighttime sleep — remained intact (Figure 1A). The single mutants showed a similar but milder phenotype, largely restricted to nighttime (Figure 1A-D). Nighttime sleep in *dArc1^-/-^* flies approached that of *dArc^-/-^*flies, but episode duration was markedly longer in the double mutant than in either single mutant, suggesting that *dArc1* and *dArc2* function synergistically to limit sleep (Figure 1C).

**Figure 1.**
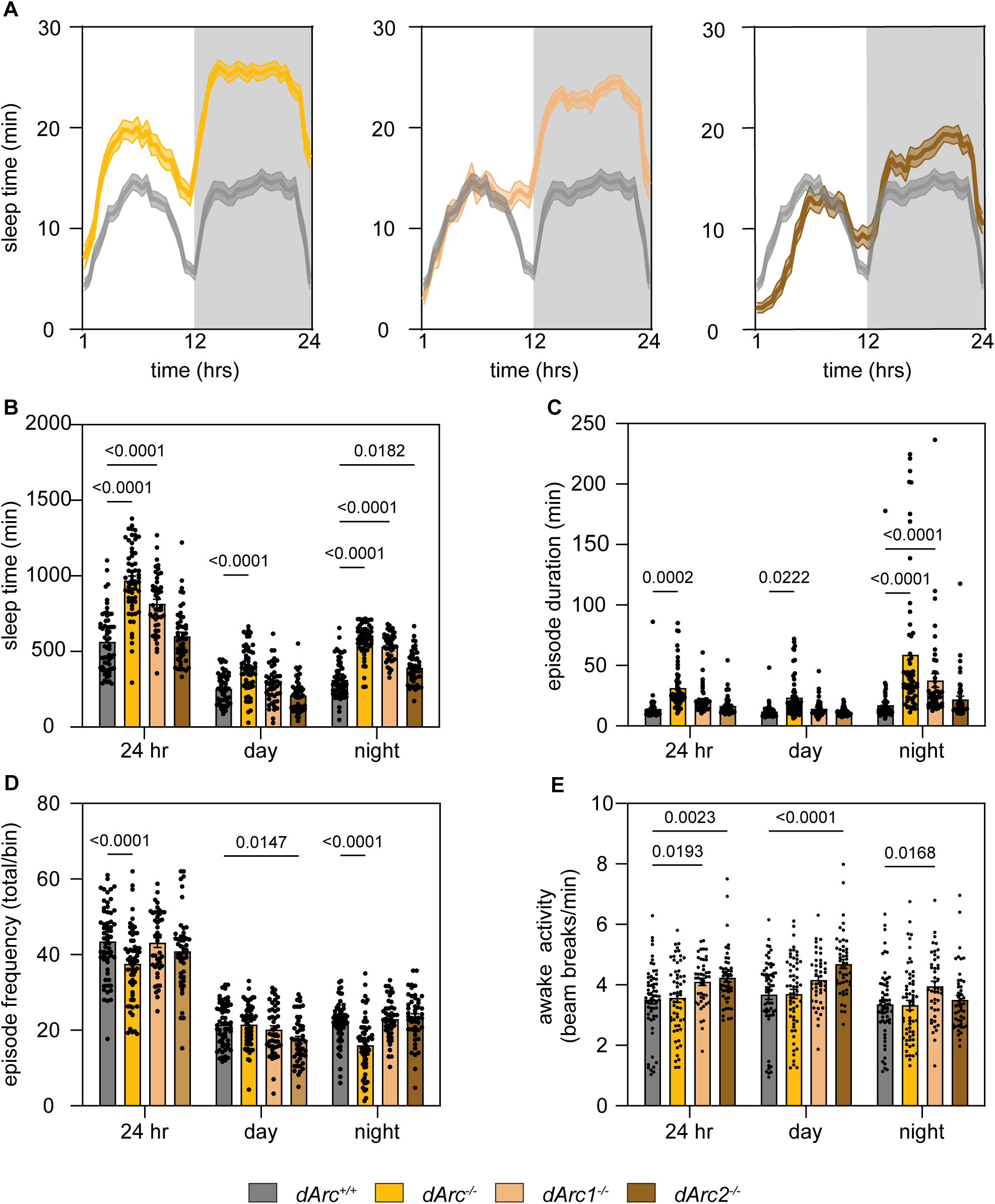
Loss of *dArc* genes increases sleep without disrupting the diurnal sleep pattern. (A) Representative sleep traces, minutes asleep per 30-minute bin, averaged across four days. (B–D) Total sleep (B), sleep episode duration (C), and sleep episode number (D), each quantified across the full day (ZT1–24), daytime (ZT1–12), and nighttime (ZT12–24). (E) Wake activity, beam breaks per waking minute, across the full day, daytime (ZT0–12), and nighttime (ZT12–24). Data are mean ± SEM. Two-way ANOVA with multiple comparisons. *dArc^+/+^*(grey, n = 60), *dArc^-/-^* (gold, n = 60), *dArc1^-/-^* (tan, n = 45), *dArc2^-/-^* (brown, n = 48). Females, 3–10 days old. See also Figures S1 and S2.

Because sleep was inferred from locomotor activity, we confirmed that these phenotypes did not reflect a locomotor defect: *dArc^-/-^* flies climbed normally^12^ and showed awake activity rates indistinguishable from *dArc^+/+^* flies (Figure 1E). These effects were present in both sexes (Figure S2A-D) but stronger in females, on which we focused all subsequent analyses.

Together, these results show that loss of *dArc* genes increases and consolidates sleep without disrupting the diurnal sleep pattern.

### Loss of *dArc* genes increases sleep depth and arousal thresholds

To determine whether the increased sleep in *dArc* mutant flies reflected deeper sleep, greater sleep pressure, or both, we applied a probabilistic analysis in which P(wake) indexes sleep depth and P(doze) indexes sleep pressure^25^. Sleep depth reflects the degree to which a fly is disengaged from external stimuli, whereas sleep pressure reflects the homeostatic drive to sleep that builds during waking and dissipates during sleep. *dArc^-/-^* flies showed a robust reduction in P(wake) across the day and night, indicating deeper sleep, while the single mutants showed reduced P(wake) only at night, consistent with their largely nighttime phenotype (Figure 2A, Figure S3A). All *dArc* mutant flies also showed a modest increase in nighttime P(doze) (Figure 2B, Figure S3B).

**Figure 2.**
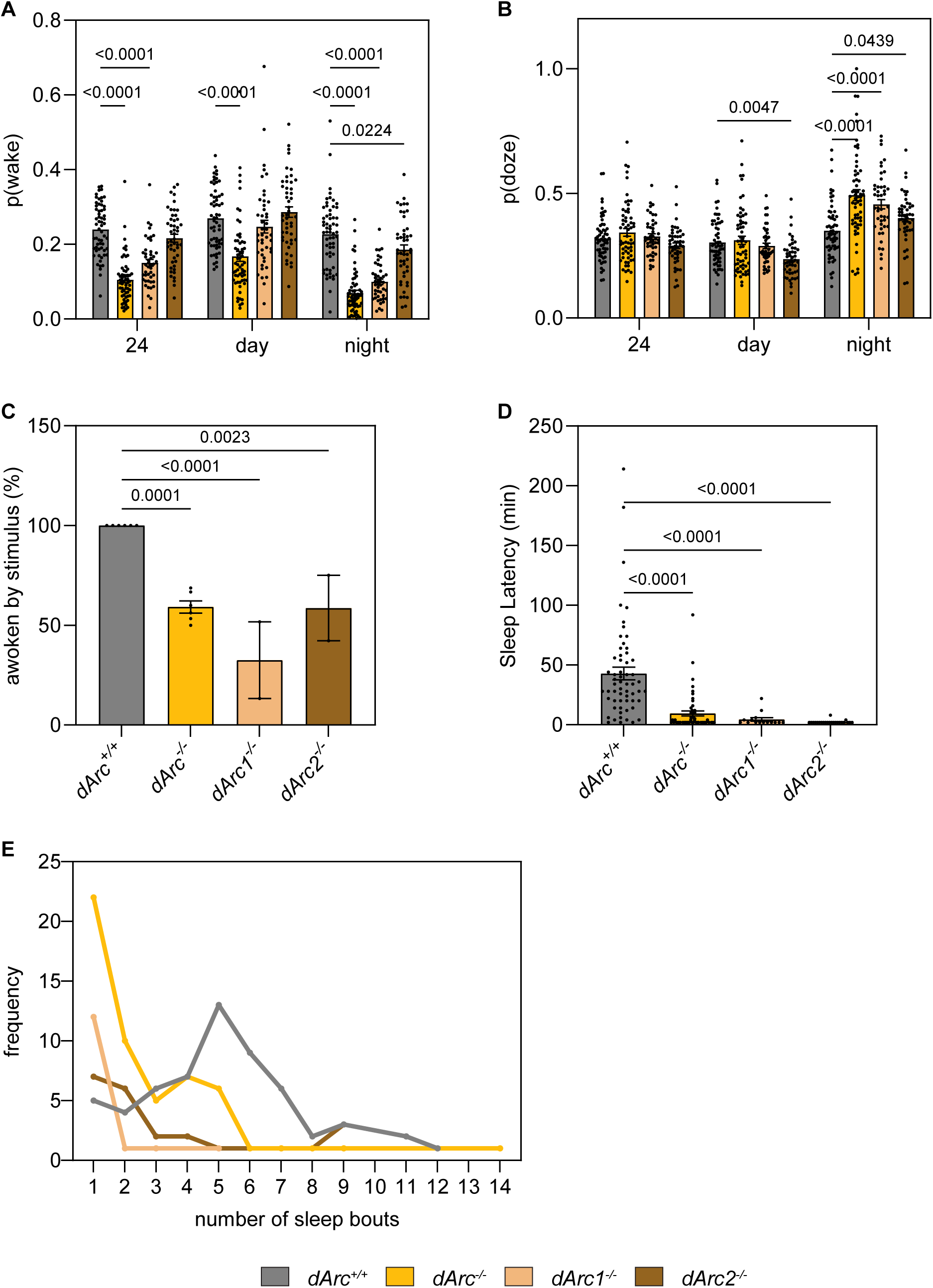
Loss of *dArc* genes increases sleep depth and arousal thresholds. (A) Average P(wake) across the full day (ZT 1-24), daytime (ZT1–12), and nighttime (ZT13–24). (B) Average P(doze) across the full day (ZT 1-24), daytime (ZT1–12), and nighttime (ZT13–24). Data are mean ± SEM. Two-way ANOVA with multiple comparisons. (A–B) *dArc^+/+^* (grey, n = 60), *dArc^-/-^* (gold, n = 60), *dArc1^-/-^* (tan, n = 45), *dArc2^-/-^* (brown, n = 48). Females, 3–10 days old. (C) Percent of flies awoken by a mechanical stimulus delivered at ZT18 for 3 seconds. Each dot represents the proportion of 32 flies that awoke. (D) Of the flies that were awoken, the time taken until the next sleep episode was recorded. (E) Of the flies that were awoken, the distribution of frequency of the number of sleep episodes occurring following mechanical stimulation. Data are mean ± SEM. One-way ANOVA with multiple comparisons. See also Figure S3.

Sleep depth is a state of sleep characterized by suppressed locomotion and reduced responsiveness to external stimuli. We therefore tested directly whether arousal thresholds were elevated in *dArc* mutant flies. A low-level mechanical stimulus delivered at ZT18, when sleep peaks, woke all *dArc^+/+^* flies but at most 59% of *dArc^-/-^* flies (Figure 2C). *dArc^-/-^* flies that did wake returned to sleep faster than *dArc^+/+^* flies (Figure 2D) and showed a longer first sleep episode after the stimulus (Figure S3C), consistent with their increased sleep depth, and the frequency of sleep episodes following the stimulus was reduced (Figure 2E). The single mutants showed similar phenotypes (Figure 2C–E). A stronger mechanical stimulus woke nearly all *dArc^-/-^* flies, although the single mutant flies remained hypoarousable (Figure S3D). These results are consistent with a prior pan-neuronal RNAi screen in which knockdown of *dArc1* produced a hypoarousable phenotype to mechanical stimulation^26^.

Because high-intensity mechanical stimulation induces a stress response that can confound arousal measurements^27^, we repeated the experiment with a light stimulus delivered at ZT18. *dArc^+/+^* and *dArc* mutant flies did not differ in their arousal to light (Figure S3E). Light and mechanical stimuli engage different upstream pathways: mechanical stimuli are detected by sensory neurons of the antenna and body wall that lie upstream of the dopaminergic system^28,29^, whereas light is detected by photoreceptors that are upstream of a distributed system of cryptochrome-expressing neurons^30,31^. That *dArc^-/-^* flies showed a defect in mechanical but not light arousal is therefore consistent with an effect on the dopaminergic system.

Together, these results show that loss of *dArc* genes deepens sleep, increasing both sleep depth and the arousal threshold in response to mechanical stimulation.

### *dArc1* is required in serotonergic neurons for setting sleep depth

We next asked which neurons require *dArc1* to regulate sleep. We previously found that dArc1 protein is detected in a small number of neurons, most of them serotonergic, and is undetectable in PAM neurons^12^; other studies, however, found that *dArc1* transcripts localize to mushroom body compartments implicated in wakefulness^19,32^. Transcriptomic data show *dArc1* transcripts across multiple neuron types, including serotonergic and dopaminergic neurons, whereas *dArc2* transcripts are barely above detection^33^. We therefore focused on *dArc1*.

*dArc1* was knocked down in specific neurons using RNA interference and the GAL4/UAS system, and knockdown was validated using RT-qPCR (Figure S4A). Knockdown in serotonergic neurons, using *tryptophan hydroxylase-GAL4*, reproduced the main phenotype observed in *dArc1^-/-^*flies: flies slept for longer periods, primarily at night, with longer sleep episodes and reduced P(wake) (Figure 3A-D, Figure S4B-D). Knockdown in PAM neurons, a subset of dopaminergic neurons that promote wakefulness, using *R58E02-GAL4*, produced a milder phenotype: flies slept for longer periods, primarily during the night, with no change in sleep episode duration or P(wake) (Figure 3E-H).

**Figure 3.**
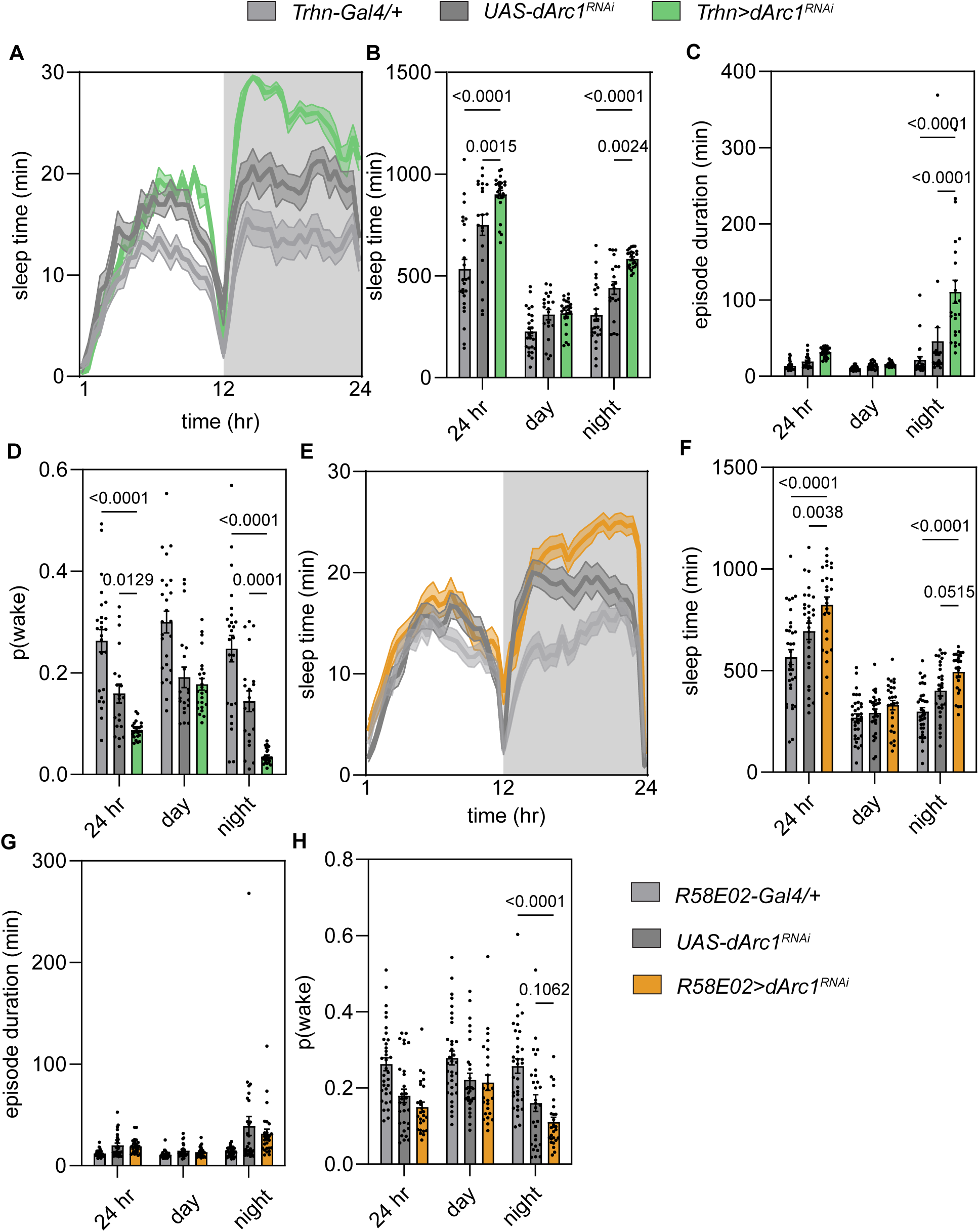
*dArc1* functions in serotonergic and PAM neurons to regulate sleep depth. (A–D) Effect of dArc1 knockdown in serotonergic neurons. (A) Representative sleep traces, minutes asleep per 30 minute bin, averaged across four days; (B) total sleep, (C) sleep episode duration, and (D) average P(wake), each across 24 hours, daytime (ZT1–12), and nighttime (ZT12–24). *Trhn-GAL4/+* (light grey, n = 25), *UAS-dArc1 RNAi/+* (grey, n = 20), *Trhn>dArc1 RNAi* (light green, n = 24). (E–H) Effect of *dArc1* knockdown in PAM neurons. (E) Representative sleep traces minutes asleep per 30 minute bin, averaged across four days; (F) total sleep, (G) sleep episode duration, and (H) average P(wake), each across 24 hours, daytime, and nighttime. *R58E02-GAL4/+* (light grey, n = 34), *UAS-dArc1 RNAi/+* (grey, n = 29), *R58E02>dArc1 RNAi* (light orange, n = 26). Data are mean ± SEM. Two-way ANOVA with multiple comparisons. Females, 3–10 days old. See also Figure S4.

Together, these results show that *dArc1* functions primarily in serotonergic neurons and to a lesser extent in PAM neurons, in both cases affecting sleep mainly at night.

### *dArc1* expression restores sleep depth in a capsid-dependent manner

We next asked whether restoring *dArc1* expression in specific neurons can rescue the sleep depth phenotype in *dArc^-/-^* flies. We expressed a *UAS-dArc1* transgene with the GAL4 lines above, all in the *dArc^-/-^* background. Restoring *dArc1* expression in serotonergic neurons produced an intermediate rescue: total sleep fell relative to *dArc^-/-^* controls but remained above *dArc^+/+^* levels, with no effect on episode duration but a full rescue of daytime sleep depth (Figure S5A-D). Expressing *dArc1* in PAM neurons produced, however, a stronger effect: daytime sleep was restored to *dArc^+/+^* levels, with episode duration and P(wake) values matching those of *dArc^+/+^* flies; nighttime sleep was partially restored (Figure 4A-D). PAM-specific expression thus affected daytime sleep most strongly.

**Figure 4.**
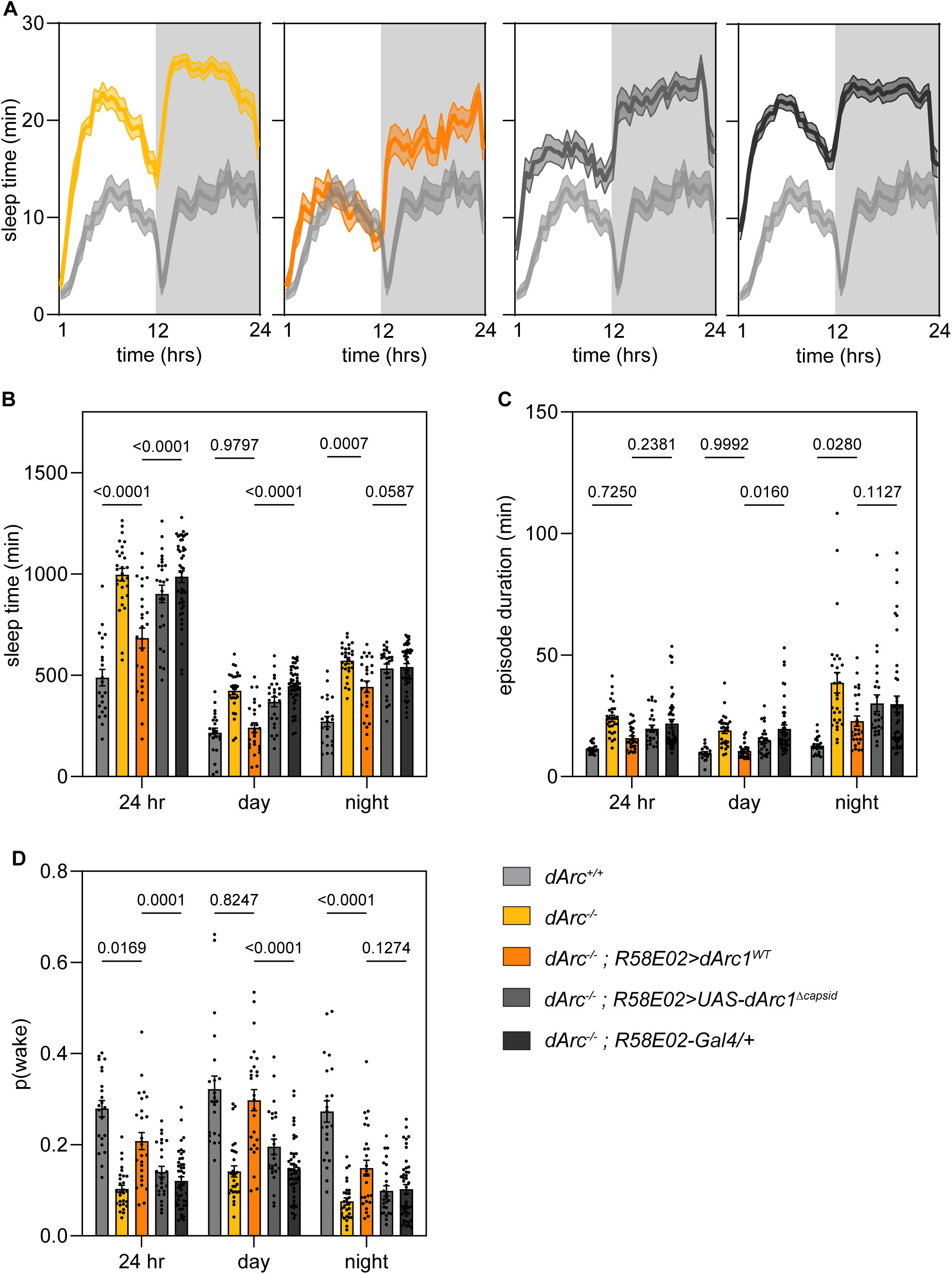
*dArc1* expression in PAM neurons rescues sleep depth in a capsid-dependent manner. (A) Representative sleep traces, minutes asleep per 30 minute bin, averaged across four days; (B) total sleep, (C) sleep episode duration, and (D) average P(wake), each across 24 hours, daytime (ZT1–12), and nighttime (ZT12–24). Data are mean ± SEM. Two-way ANOVA with multiple comparisons. *dArc^+/+^*(light grey, n = 22), *dArc^-/-^* (gold, n = 28), *dArc^-/-^; R58E02>dArc1* (orange, n = 26), *dArc^-/-^; R58E02> dArc1^Δcapsid^* (grey, n = 25), *dArc^-/-^; R58E02/+* (dark grey, n = 44). Females, 3–10 days old. See also Figure S5.

To test whether *dArc1* function in serotonergic and PAM neurons required capsid formation, we used a *dArc1* variant bearing two mutations that prevent capsid assembly^12^ (*dArc1^Δcapsid^*). Expressing *dArc1^Δcapsid^* in serotonergic neurons of *dArc^-/-^* flies failed to restore sleep to *dArc^+/+^* levels (Figure S5A-D). Likewise, expressing it in PAM neurons did not significantly restore sleep amount, episode duration, or P(wake), though trend-level differences were present for all three (Cohen’s d = 0.71, 0.51, and 0.67; Figure 4B-D). These trends may reflect residual activity of *dArc1^Δcapsid^*. In either case, capsid formation was required for full rescue.

Together, these results show that *dArc1* expression in either serotonergic or PAM neurons restores daytime sleep depth, with the stronger effect in PAM neurons, and that capsid formation is required in both cases.

### *dArc1* in Sas-expressing cells and Ptp10D in PAM neurons are required for sleep depth

These results, together with prior work^12^, suggest that serotonergic neurons deliver dArc1 capsids to PAM neurons. A candidate mechanism for this delivery has been previously described: Sas, a cell surface protein present on extracellular vesicles, binds dArc1 through its cytoplasmic domain and targets those vesicles to cells expressing its receptor Ptp10D, facilitating delivery of dArc1 capsids into recipient cells^13^. Both transcripts are expressed in the adult brain^33,34^. We therefore tested whether this system mediated dArc1 signaling during sleep.

Knocking down *dArc1* in Sas-expressing cells did not change total sleep but increased nighttime episode duration and reduced nighttime P(wake) (Figure S6A-D), indicating that *dArc1* is required in Sas-expressing cells for sleep depth. Knocking down *Ptp10D* in PAM neurons, using *R58E02-GAL4*, increased nighttime sleep and episode duration and reduced nighttime P(wake) (Figure 5A-D), reproducing features of the phenotype observed in *dArc^-/-^* flies.

**Figure 5.**
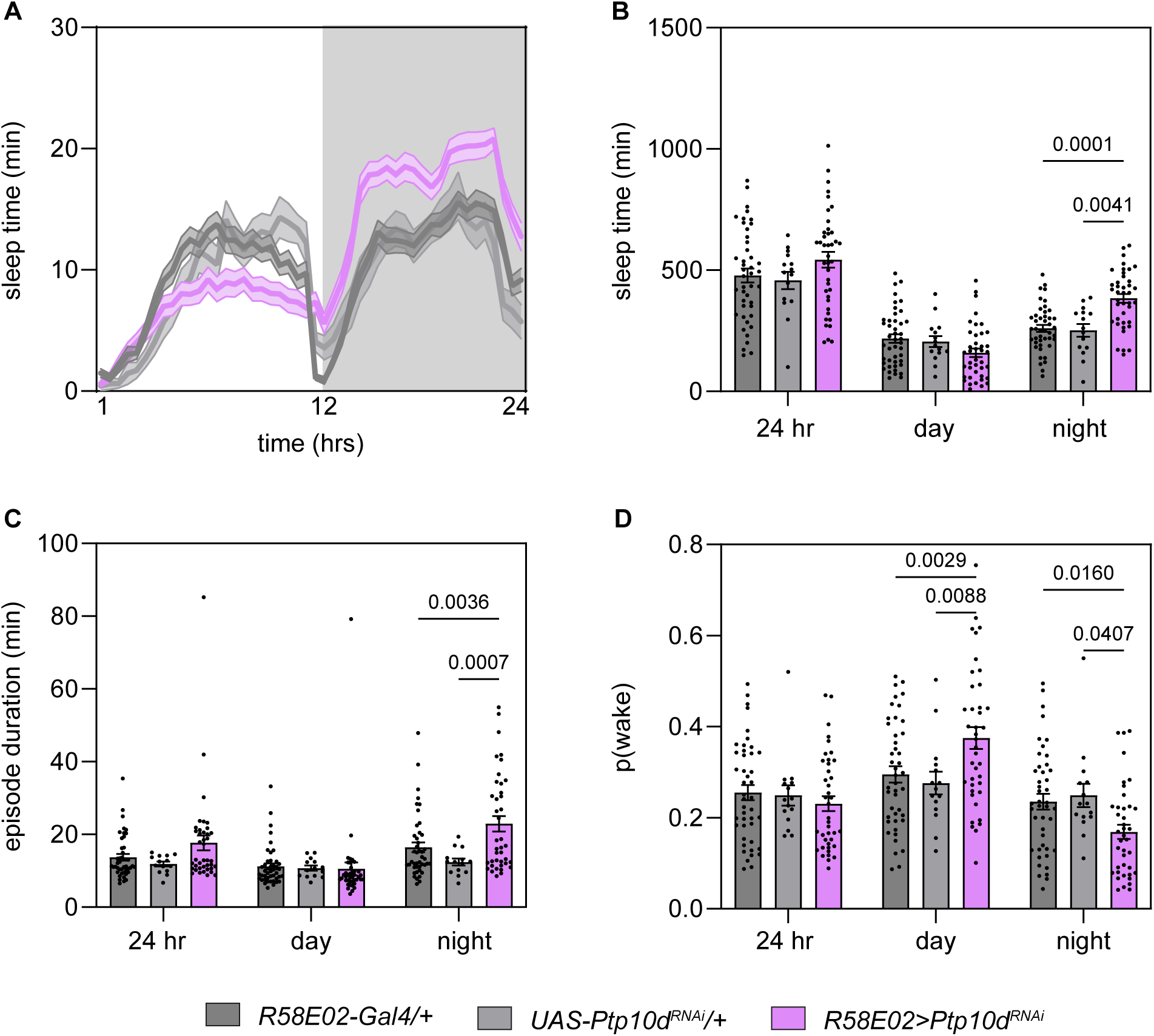
Ptp10D is required in PAM neurons to control sleep depth. (A) Representative sleep traces, minutes asleep per 30-minute bin, averaged across four days; (B) total sleep, (C) sleep episode duration, and (D) average P(wake), each across 24 hours, daytime (ZT1–12), and nighttime (ZT12–24). Data are mean ± SEM. Two-way ANOVA with multiple comparisons. R58E02-Gal4/+ (grey, n = 44), UAS-Ptp10DRNAi (light grey, n = 15) R58E02>Ptp10DRNAi (pink, n = 40). Females, 3–10 days old. See also Figure S6.

Together, these results support a model in which Sas-tagged extracellular vesicles deliver dArc1 capsids to Ptp10D-expressing PAM neurons to regulate sleep depth.

## DISCUSSION

The results presented here show that *dArc1* sets sleep depth through capsid-mediated signaling between serotonergic and PAM neurons, and that this signaling most likely involves the Sas-Ptp10D machinery. Arc-dependent capsid signaling therefore operates as a mechanism that couples two modulatory neuronal populations to set sleep depth.

The sleep phenotype in *dArc^-/-^* flies is an increase in sleep depth rather than a decrease in the drive to wake. Two pieces of evidence support this. First, *dArc^-/-^* flies show a modality-specific arousal phenotype that points to the dopaminergic system: flies are hypoaroused to weak mechanical stimuli, which are relayed through the dopaminergic system^28,29^, but respond to light as readily as control flies, arousal that is relayed through the cryptochrome-expressing system^30,31^. Second, P(wake), the probability that a sleeping fly resumes activity, is reduced in *dArc^-/-^* flies. Combined with the finding that *dArc1* functions in PAM neurons, an arousal-promoting population, these results suggest that the consequence of *dArc* loss is not a shift in wake drive but a lowering of the arousal-promoting output of the PAM neurons, which deepens sleep while leaving its diurnal sleep pattern intact. Loss of *dArc1* affected sleep mainly at night, whereas its restoration acted mainly during the day. This difference may reflect the temporal regulation of *dArc1*: endogenous expression may vary across the day and night, whereas the GAL4/UAS system used for rescue drives expression without such regulation. Whether *dArc1* expression is regulated across the day and night remains to be tested.

How *dArc1* signals from serotonergic to PAM neurons most likely involves Sas and Ptp10D, a ligand-receptor pair first described in axon guidance^35^ and later shown to transfer dArc1 capsids in non-neuronal tissues^13^. Knockdown of *dArc1* in Sas-expressing cells, or of *Ptp10D* in PAM neurons, reproduced the deeper, more consolidated nighttime sleep and reduced P(wake) of *dArc^-/-^* flies, indicating that this delivery system also operates in the adult brain. Expressing *dArc1* directly in PAM neurons supplies the gene without transfer from serotonergic neurons, and this manipulation restored sleep depth. The capsid-deficient variant expressed in these same neurons did not fully restore sleep depth, indicating that capsid assembly is required for *dArc1* to function within PAM neurons and not only for its transfer between serotonergic and PAM neurons. The most parsimonious interpretation is that the capsid does more than transfer cargo into PAM neurons: it may be required for a further signaling step, perhaps transfer from PAM neurons to downstream targets. Identifying these targets, and the RNA cargo the capsid carries, will be important goals for future work.

The capsid-forming property of Arc genes, retained independently in insects and tetrapods^3,4^, distinguishes this signal from a neuropeptide or other diffusible modulator: the capsid packages RNA^6,7^, including its own mRNA^3,4,6,7^, and transfers it into the receiving neuron^4^, such that the signal carries translatable cargo rather than acting only at a surface receptor. It is conceivable that this mode of signaling is suited to setting and maintaining a state rather than triggering a rapid one: assembling, transferring, and acting within the receiving neuron over minutes to hours, more slowly than synaptic transmission, such that its influence accumulates gradually. Consistent with this, dArc1 sets the depth of sleep, a maintained state, over a timescale of hours. Serotonergic and dopaminergic neurons integrate internal state to tune behavior to the needs of an animal, and because they also control behaviors on faster timescales, examining dArc signaling between them, and across species, could test whether the capsid acts generally as a slow, accumulating signal. Consistent with a conserved role, *Arc* knockout mice show an attenuated homeostatic response to sleep deprivation: they lack the rebound sleep that usually follow deprivation, a hallmark of the sleep homeostat, and they spend more time rapid eye movement sleep^36^. *Arc* expression itself increases after sleep deprivation in wild-type mice^36^, consistent with a role in tracking sleep need. Together with our finding that *dArc1* sets sleep depth in *Drosophila*, these observations suggest that the involvement of Arc in sleep regulation is conserved across species.

## ACKNOWLEDGMENTS

We thank members of the Caron laboratory for comments on the manuscript and discussions of the project. We are grateful to Drs. Adrian Rothenfluh, Kent Golic, and Michael Werner for their wisdom and thoughtful discussions throughout this project. We want to acknowledge Adam Lin and Sylvia Yang for preparation of the standard cornmeal agar medium; we also want to acknowledge Hayley Smihula, Ashley Platt, Miles Jacob, and Lia Beatty for providing lab management during the project. This work was initially funded by a Seed Grant awarded by the Neuroscience Initiative at the University of Utah to S.J.C.C. and J.D.S.; the project has been funded thereafter by grants from the National Institute for Neurological Disorders and Stroke (R01 NS 106018, R01 NS 107790 and R01 NS 115716) and the National Science Foundation (IOS 2042397). Further financial support was provided by a Genetic Training Grant (T32GM141848, A.R.B), the Undergraduate Research Opportunities Program (K.D.), a Biology Research Scholar Award (K.D.), a Parent Fund Scholarship (K.D.), the Chan Zuckerberg Initiative (Ben Barres Early Acceleration Award; J.D.S.), the Jon M. Huntsman Presidential Endowed Chair Fund (J.D.S.) and the George S. and Dolores Doré Eccles Foundation (S.J.C.C.).

## AUTHOR CONTRIBUTIONS

A.R.B., J.D.S., and S.J.C.C. conceived the project. A.R.B. and S.J.C.C. wrote the manuscript with input from all other authors. A.R.B. performed and analyzed the behavioral assays and validated the *dArc* mutants and RNAi knockdown lines. K.D. performed behavioral assays. S.B. generated the *dArc* mutant lines. J.D.S. and S.J.C.C. supervised the project. A.R.B., K.D., J.D.S., and S.J.C.C. acquired funding.

## RESOURCE AVAILABILITY

### Lead contact

Further information and requests for resources and reagents should be directed to and will be fulfilled by the lead contact, Sophie Caron.

### Materials availability

Plasmids and fly lines generated in this study are available upon request.

### Data and code availability

All data reported in this paper will be shared by the lead contact upon request. This paper does not report original code. Any additional information required to reanalyze the data reported in this paper is available from the lead contact upon request.

## STAR METHODS

### Key Resources Table

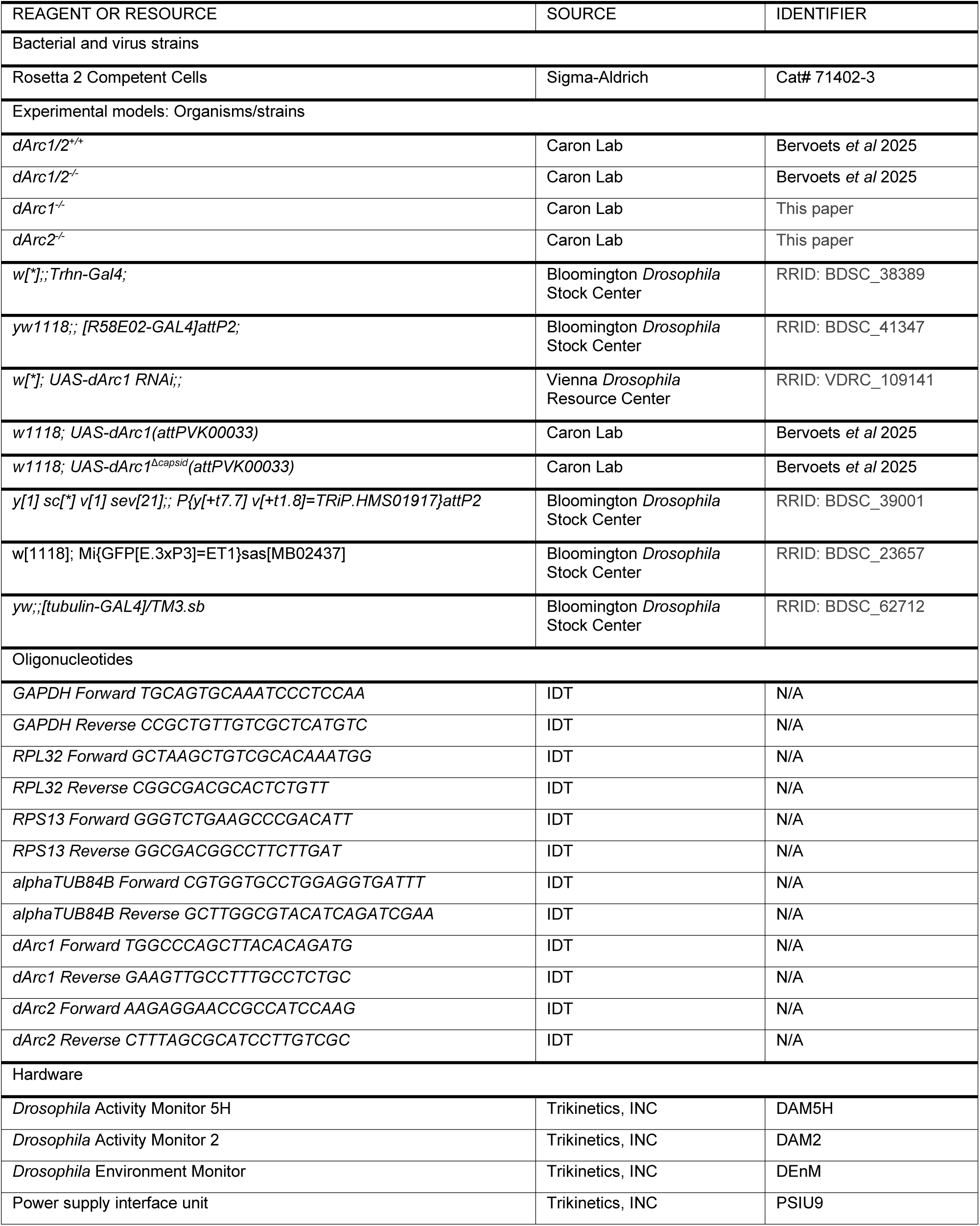
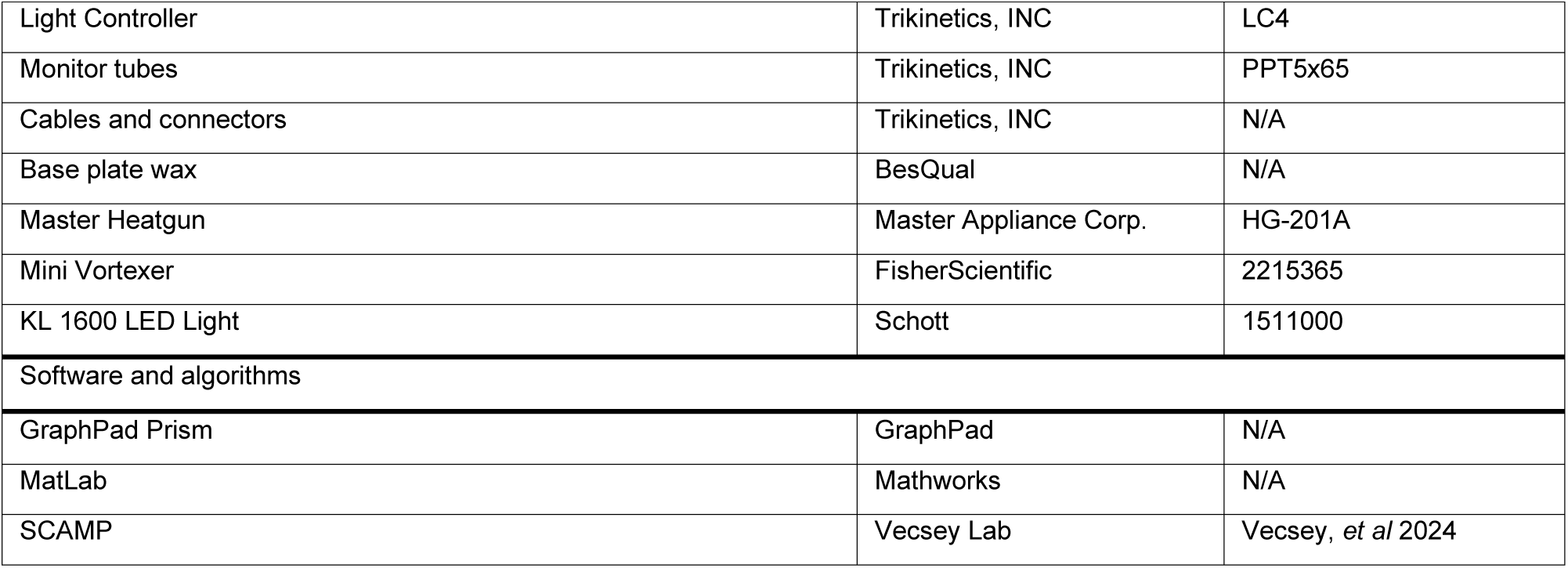

### EXPERIMENTAL MODELS AND STUDY PARTICIPANTS

#### Fly stocks and husbandry

Flies (*Drosophila melanogaster*) were reared on standard cornmeal agar medium (cornmeal 50.8 g/L, agar 9.3 g/L, yeast 15 g/L, Karo light corn syrup 12.5 mL/L, malt 37.03 mL/L, molasses 18.52 mL/L, Tegosept 8.33 mL/L, propionic acid 4.16 mL/L) in a controlled environmental chamber (Percival Scientific, Perry, IA) maintained at 25°C and 60% relative humidity under a 12-hour light, 12-hour dark cycle. Crosses were reared under the same conditions, except for RNAi crosses, which were reared at 28°C to maximize knockdown efficiency. Sex, age, and number of flies are given in the figure legends. All fly strains used in this study are listed in the Key Resources Table.

#### Microbe strains

For cloning, NEB Stable Competent E. coli were grown on LB agar and LB medium with ampicillin at 37◦ C. For protein expression, Rosetta 2 BL21 competent E. coli were grown overnight at 37◦ C in LB medium supplemented with ampicillin and chloramphenicol. Large-scale ZY auto-induction medium was inoculated with the starter culture and grown at 37◦ C until a 600 nm optic density of 0.6-0.8 with shaking at 160 rpm, then shifted to 18◦ C with shaking at 180 rpm.

### METHOD DETAILS

#### Generation of *dArc1^-/-^* and *dArc2^-/-^* flies

*dArc1^-/-^* and *dArc2^-/-^* flies were generated using CRISPR/Cas9-mediated genomic engineering, following an established protocol^37,38^. Briefly, four guide RNAs (gRNAs) were designed to target various regions of dArc1 and dArc2 (gRNA1-4) and were evaluated for specificity and off-target effects using the CRISPR Target Finder tool (http://targetfinder.flycrispr.neuro.brown.edu/). These gRNA sequences were cloned into the pU6-BbsI-chiRNA plasmid and sequence-verified. Homology arms of approximately 1 kb flanking the Cas9 cut sites were synthesized as gene blocks (Integrated DNA Technologies, Coralville, IA), cloned into the pHD-DsRed-attP plasmid, and sequence-verified. The gRNAs and homology repair plasmids were co-injected into vas-Cas9 transgenic embryos (BestGene Inc., Chino Hills, CA). Adult flies were backcrossed to *w1118* flies. Mutant chromosomes were identified using the DsRed marker, which was under the 3xP3 promoter and therefore visible in adult eyes, and isolated using the balancer chromosome CyO. The resulting *dArc1^-/-;DsRed^* and *dArc2^-/-;DsRed^*flies were crossed to flies expressing Cre recombinase to excise the DsRed marker, yielding the *dArc1^-/-^* and *dArc2^-/-^* flies used throughout this study. Both lines were validated by PCR using primers flanking the targeted region. The control line, *dArc^+/+^,* was established by isolating the second chromosome from vas-Cas9 flies.

#### Reverse-transcription quantitative polymerase chain reaction (RT-qPCR)

RT-qPCR was performed as previously described^39^. Briefly, total RNA was extracted from three biological replicates per condition using a standard TRIzol (Qiagen, Hilden, Germany) and chloroform extraction protocol, followed by isopropanol precipitation and 75% ethanol washes. The precipitated RNA was treated with recombinant DNase I (Roche, Basel, Switzerland). Single-stranded cDNA was synthesized using the High-Capacity cDNA Reverse Transcription Kit (Applied Biosystems, Thermo Fisher Scientific, Waltham, MA). Gene expression levels were determined using PowerUp SYBR Green Master Mix (Applied Biosystems, Thermo Fisher Scientific, Waltham, MA) and measured on a QuantStudio 3 real-time PCR system (Applied Biosystems, Thermo Fisher Scientific, Waltham, MA). Expression was normalized using stable housekeeping genes: gene stability was first assessed across samples, and the most stable genes were selected to normalize target gene expression. Primer sequences are listed in the Key Resources Table.

#### Locomotor recording and sleep analysis

Locomotor behavior was recorded using the *Drosophila* Activity Monitor system 5 (DAM5H, Trikinetics, Waltham, MA). On day zero, three- to seven-day-old flies were collected between ZT2 and ZT4 under CO_2_ anesthesia and loaded into individual glass tubes containing standard cornmeal agar medium (recipe described above), allowing one day of acclimation before recording. Each tube was capped with dental wax at one end and sealed with a small piece of laboratory tissue at the other. Tubes were loaded into DAM5H boards. The DAM5H boards use four independent infrared beams per fly to track activity. Data collection began at ZT1 on day one and continued for four days. Recordings were made in an environmental chamber (Percival Scientific, Perry, IA) under a 12-hour light, 12-hour dark cycle at 25°C and 60% relative humidity.

Sleep was defined as any period of inactivity lasting five minutes or longer as is standard in the field^22,23^. Activity data were analyzed using the MATLAB-based program SCAMP^24^, from which total sleep, sleep episode duration, and sleep episode number were extracted for the full 24-hour period and separately for the 12-hour day and 12-hour night. Wake activity was calculated as the number of beam breaks *per* waking minute. Sleep quality was assessed by computing P(wake) and P(doze) as established by a previous study^25^. P(wake), the probability that an inactive fly becomes active in the following minute, indexes sleep depth, with lower values reflecting deeper sleep; P(doze), the probability that an active fly becomes inactive in the following minute, indexes sleep pressure, with higher values reflecting a greater drive to sleep. Both metrics were averaged across the 24-hour period and separately for the day and night.

#### Arousal assays

Mechanical arousal: Locomotor behavior was recorded as described above, using the Drosophila Activity Monitor system 2 (DAM2, Trikinetics, Waltham, MA), which uses a single infrared beam per fly to track activity. Three- to seven-day-old female flies were collected between ZT2 and ZT4 under CO_2_ anesthesia and loaded into individual glass tubes containing standard cornmeal agar medium, prepared as described for the locomotor recording assay. Tubes were loaded into DAM2 boards, secured with rubber bands, and mounted on a modified VortexGenie2 (Scientific Industries, Bohemia, NY). Data collection began at ZT1 on day one, and a three second mechanical stimulus was delivered at ZT18 on night one. Recordings were made in an environmental chamber (Percival Scientific, Perry, IA) under a 12-hour light, 12-hour dark cycle at 25°C and 60% relative humidity. Arousal was scored manually in a custom Excel spreadsheet: a fly was considered asleep if it showed six minutes or more of inactivity before the stimulus, and aroused if it showed activity in the two minutes following the stimulus.

Light arousal: Locomotor behavior was recorded using the Drosophila Activity Monitor system (DAM5H, Trikinetics, Waltham, MA) under the same collection, loading, and housing conditions as the mechanical arousal assay. Data collection began at ZT1 on day one, and at ZT18 on night one the incubator lights were turned on for two minutes. Arousal was analyzed using the MATLAB-based program SCAMP^13^.

### QUANTIFICATION AND STATISTICAL ANALYSIS

Measurements were taken from independent samples. Statistical analyses were performed in GraphPad Prism 10.4.1 (GraphPad Software, Boston, MA). Sample sizes were determined by power analyses on pilot experiments. Alpha was set at 0.05, and significance was assigned to p-values below 0.05. Numerical p-values are reported in the figure legends. The statistical test, sample size, and error bars for each experiment are given in the corresponding figure legend.

## Supplemental Information

***dArc1* downregulates sleep depth through capsid-mediated signaling to dopaminergic neurons**

**Figure S1.**
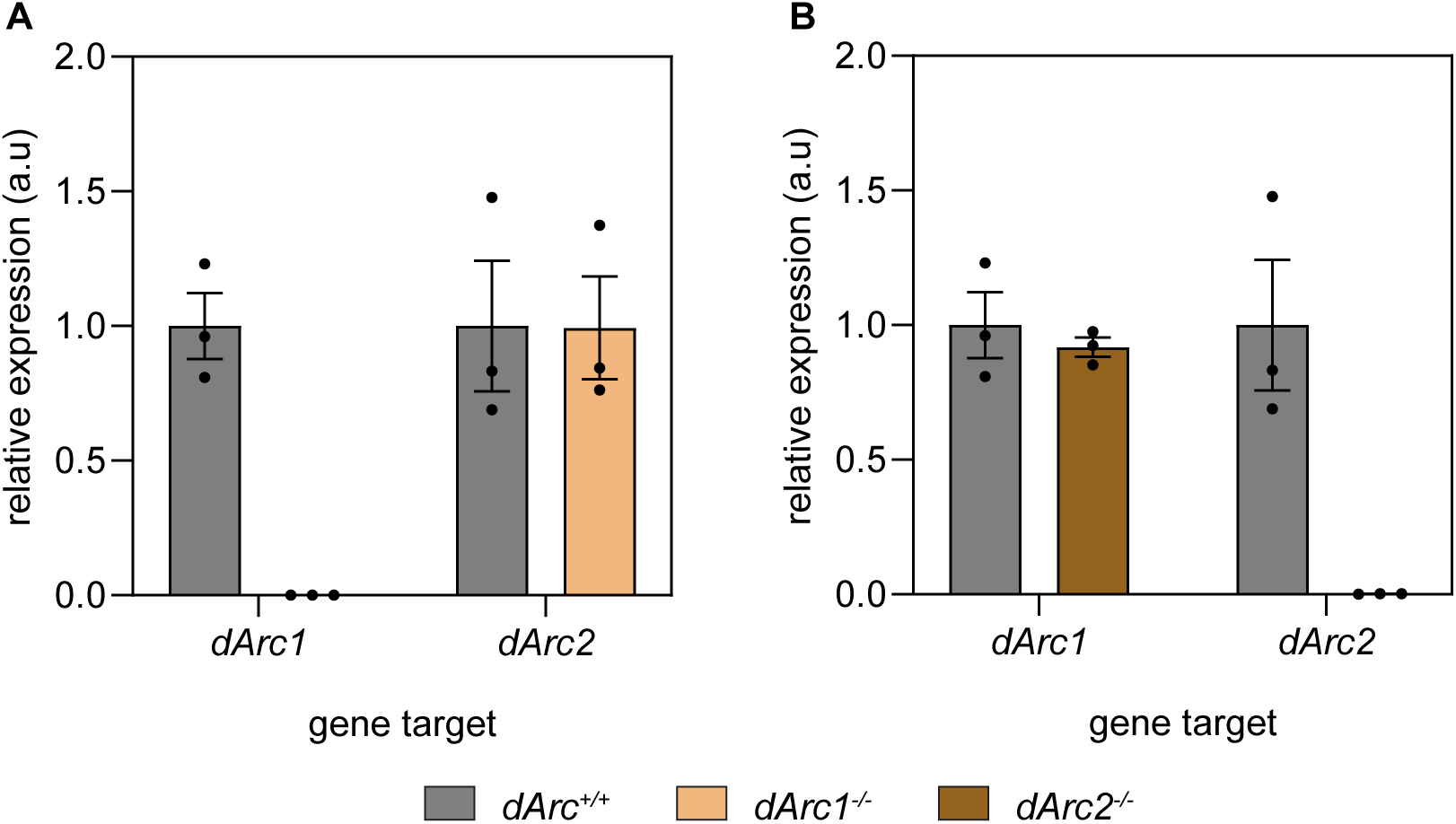
Validation of *dArc1^-/-^* and *dArc2^-/-^* mutations. (A) Relative mRNA expression of *dArc1* and *dArc2* from *dArc^+/+^* and *dArc1^-/-^* assessed via RT-qPCR. (B) Relative mRNA expression of *dArc1* and *dArc2* from *dArc^+/+^* and *dArc2^-/-^* assessed via RT-qPCR. Each dot represents an independent sample of 5-10 flies of mixed sex.

**Figure S2.**
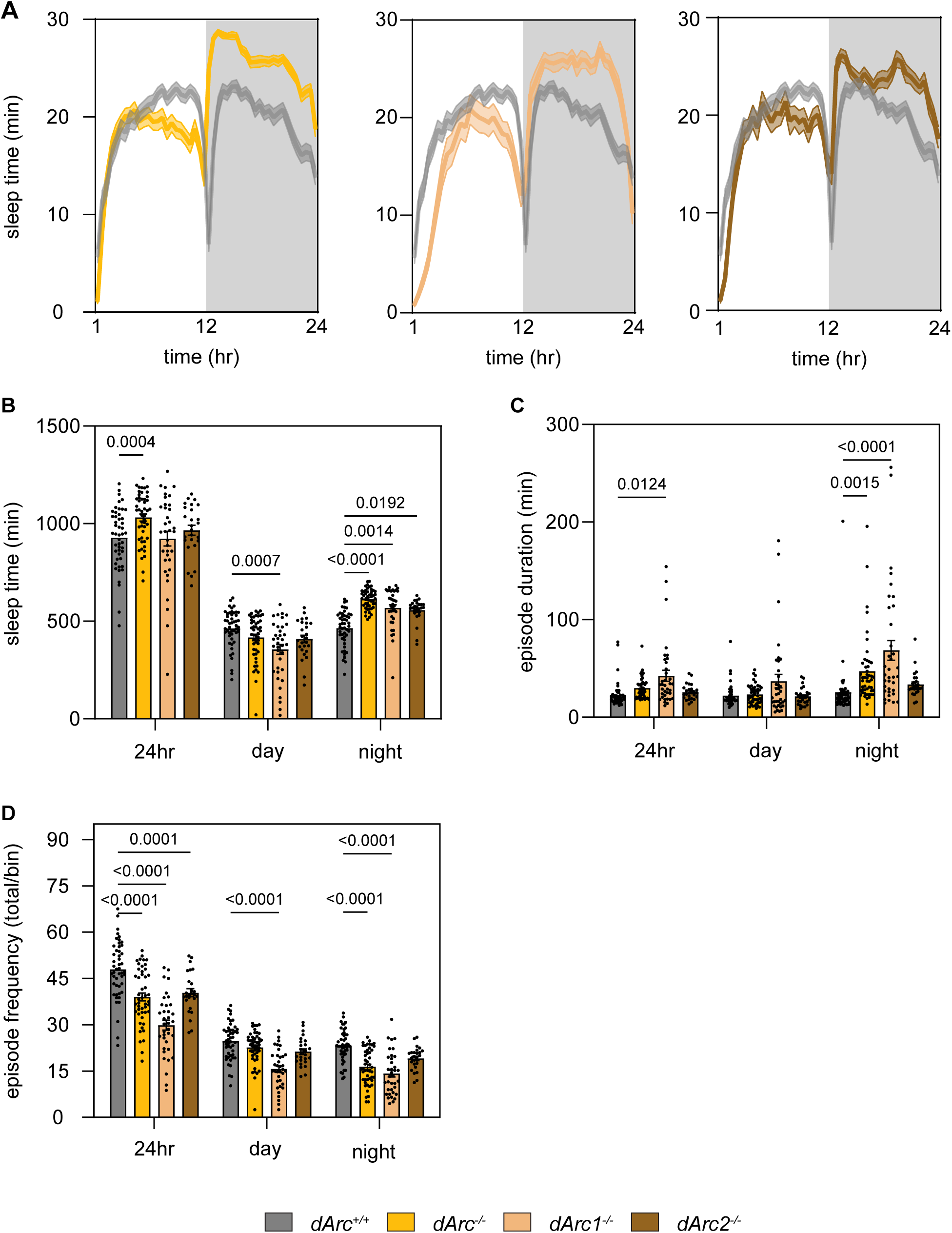
Loss of *dArc* genes increases sleep without disrupting the diurnal sleep pattern. (A) Representative sleep traces, minutes asleep per 30-minute bin, averaged across four days. (B–D) Total sleep (B), sleep episode duration (C), and sleep episode number (D), each quantified across the full day (ZT1–24), daytime (ZT1–12), and nighttime (ZT13–24). (E) Wake activity, beam breaks per waking minute, across the full day, daytime (ZT0–12), and nighttime (ZT12–24). Data are mean ± SEM. \**p* < 0.05, \*\**p* < 0.01, \*\*\**p* < 0.001, \*\*\*\**p* < 0.0001, two-way ANOVA with multiple comparisons. *dArc^+/+^*(grey, n = 48), *dArc^-/-^* (gold, n = 48), *dArc1^-/-^*(tan, n = 36), *dArc2^-/-^* (brown, n = 26). Males, 3–10 days old.

**Figure S3.**
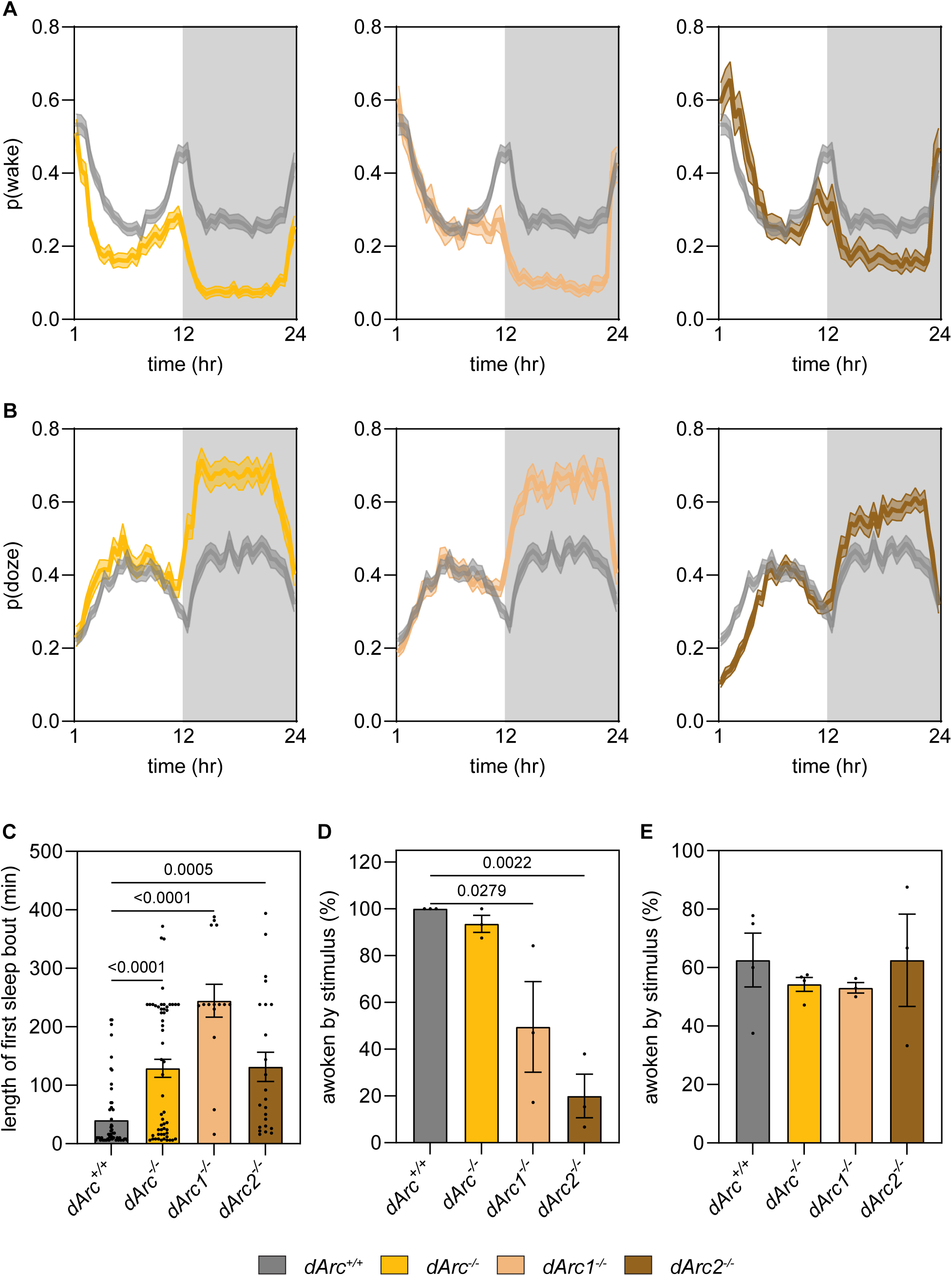
Loss of *dArc* genes increases sleep depth and arousal thresholds. (A) Representative p(wake) trace, the probability of transitioning from sleep to wakefulness, binned per 30-minute interval and averaged across four days. (B) Representative p(doze) trace, the probability of transitioning from wakefulness to sleep, binned per 30-minute interval and averaged across four days. (C) Of the flies that were awoken, the length of the first sleep episode following arousal was recorded. (D) Percent of flies awoken by a high-intensity mechanical stimulation delivered at ZT18 for 3 seconds. Each dot represents the proportion of 32 flies that awoke. (E) Percent of flies awoken by a LED stimulus delivered at ZT18 for 2 minutes. Each dot represents the proportion of 32 flies that awoke. Data are mean ± SEM. \**p* < 0.05, \*\**p* < 0.01, \*\*\**p* < 0.001, \*\*\*\**p* < 0.0001, one-way ANOVA with multiple comparisons. Females, 3–10 days old.

**Figure S4.**
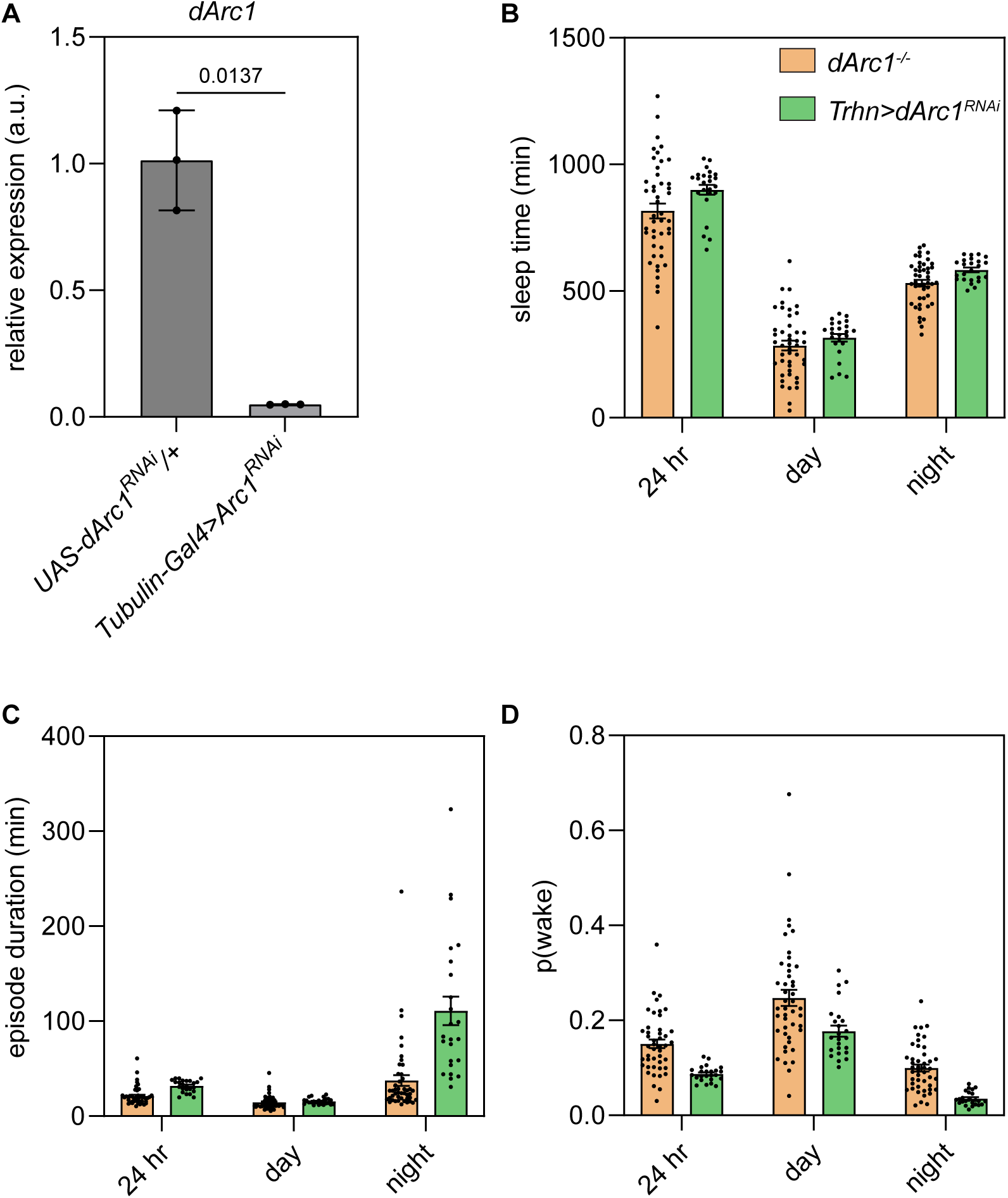
*dArc1* is necessary in serotonin and PAM dopaminergic neurons to regulate sleep depth. (A) Validation of *dArc1* knockdown via RNA interference. Expression of *UAS-dArc1^RNAi^* in all cells results in significant reduction in detectable *dArc1* mRNA. (B-D) Qualitative comparison of sleep behavior between *dArc1^-/-^* from Figure 1 and serotonin specific knockdown of *dArc1* from Figure 3. (B) total sleep, (C) sleep episode duration, and (D) average p(wake), each across 24 hours, daytime (ZT1–12), and nighttime (ZT13–24).

**Figure S5.**
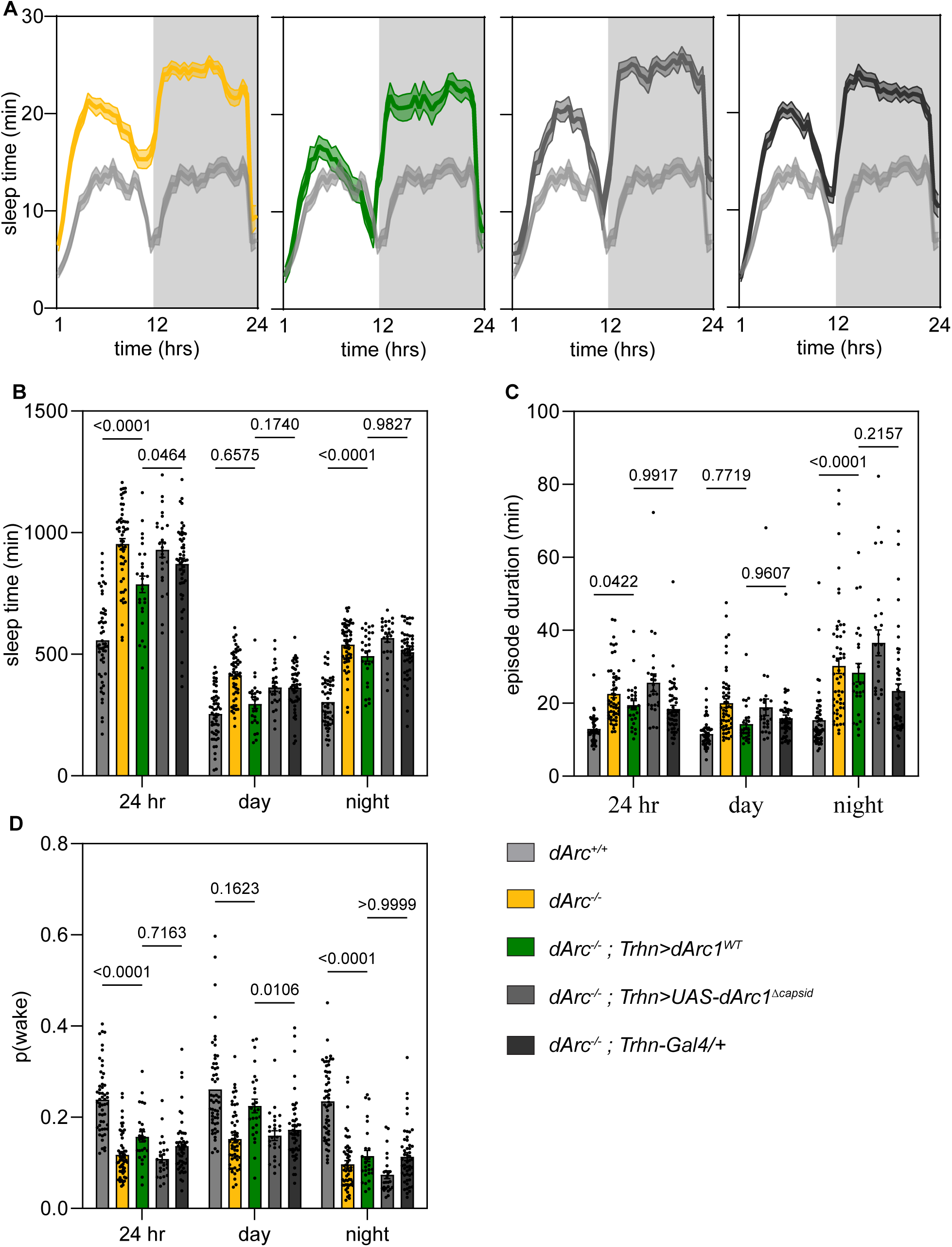
*dArc1* expression in serotonin neurons does not rescue sleep depth. (A) Representative sleep traces, minutes asleep per 30 minute bin, averaged across four days; (B) total sleep, (C) sleep episode duration, and (D) average p(wake), each across 24 hours, daytime (ZT1–12), and nighttime (ZT13–24). Data are mean ± SEM. \**p* < 0.05, \*\**p* < 0.01, \*\*\**p* < 0.001, \*\*\*\**p* < 0.0001, two-way ANOVA with multiple comparisons. *dArc^+/+^* (light grey, n = 53), *dArc^-/-^*(gold, n = 53), *dArc^-/-^; Trhn>dArc1^WT^* (green, n = 26), *dArc^-/-^; Trhn>dArc1^Δcapsid^* (grey, n = 25), *dArc^-/-^; Trhn/+* (dark grey, n = 50). Females, 3–10 days old.

**Figure S6.**
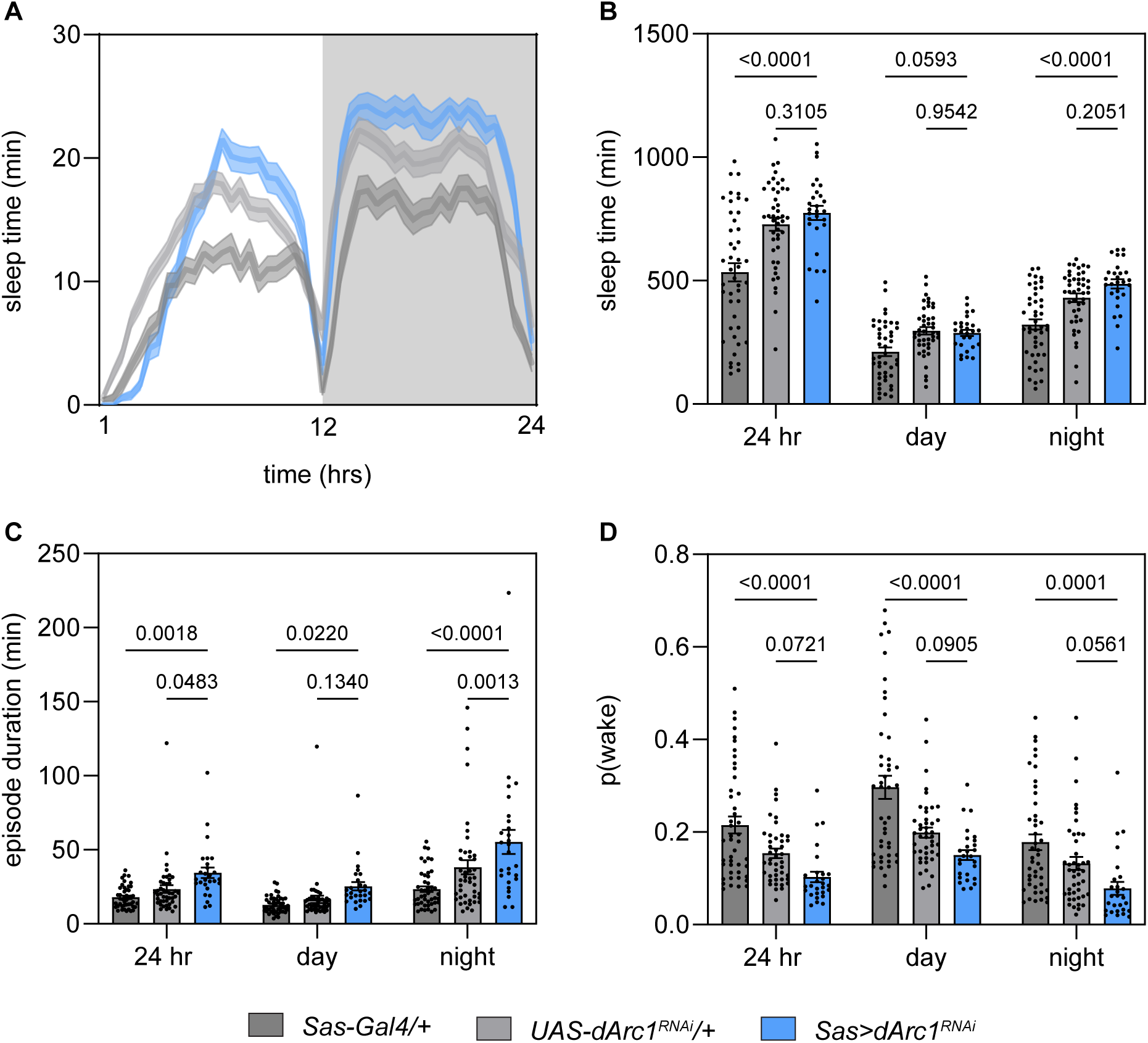
Knockdown of *dArc1* in Sas-Gal4 cells reproduces aspects of the *dArc1^-/-^* mutant. (A) Representative sleep traces, minutes asleep per 30 minute bin, averaged across four days; (B) total sleep, (C) sleep episode duration, and (D) average p(wake), each across 24 hours, daytime (ZT1–12), and nighttime (ZT13–24). Data are mean ± SEM. \**p* < 0.05, \*\**p* < 0.01, \*\*\**p* < 0.001, \*\*\*\**p* < 0.0001, two-way ANOVA with multiple comparisons. *Sas-Gal4/+* (grey, n = 45), *UAS-dArc1^RNAi^* (light grey, n = 44) *Sas>dArc1^RNAi^* (blue, n = 27). Females, 3–10 days old.

## REFERENCES

1. Zhang W, et al. Structural basis of Arc binding to synaptic proteins: implications for cognitive disease. Neuron 2015;86, 490–500. 10.1016/j.neuron.2015.03.030

2. Myrum C, et al. Arc is a flexible modular protein capable of reversible self oligomerization. Biochem Jour. 2015;1,145–158. 10.1042/BJ20141446

3. Ashley J, Cordy B, Lucia D, Fradkin LG, Budnik V, Thomson T. Retrovirus-like gag protein Arc1 binds RNA and traffics across synaptic boutons. Cell. 2018;172(1-2):262–274.e11. doi:10.1016/j.cell.2017.12.022

4. Pastuzyn, ED. et al. The neuronal gene *Arc* encodes a repurposed retrotransposon gag protein that mediates intercellular RNA transfer. Cell. 2018;172(1-2), 275-288. 10.1016%2Fj.cell.2017.12.024

5. Sullivan KR, Ravens A, Walker AC, Shepherd JD. “Arc – A viral vector of memory and synaptic plasticity.” Cur. Opi. Neurobio.. 2025;91:102979. doi:10.1016/j.conb.2025.102979

6. Zinter M, Xiao C, M’Angale P, et al. Arc capsids facilitate the transfer of muscleblind. bioRxiv. Published online March 2026.

7. Xiao C, M’Angale G, Wang S, Lemieux A, and Thompson T. Idenitifying new players in structural synaptic plasticity through *dArc1* interrogation. iScience. 2023;26,108048. 10.1016/j.isci.2023.108048.

8. Hantak MP, Einstein J, Kearns RB, and Shepherd JD. Intercellular communication in the nervous system goes viral. Trends Neurosci. 2021;44(4) 248–259. 10.1016/j.tins.2020.12.003.

9. Mattaliano MD, Montana ES, Parisky KM, Littleton JT, Griffith LC. The Drosophila ARC homolog regulates behavioral responses to starvation. Mol. Cell. Neurosci. 2007;36(2):211–221. doi:10.1016/j.mcn.2007.06.008

10. Mosher J, Zhang W, Blumhagen RZ, et al. Coordination between Drosophila Arc1 and a specific population of brain neurons regulates organismal fat. Dev. Bio. 2015;405(2):280–290. doi:10.1016/j.ydbio.2015.07.021

11. Awata H, Takakura M, Kimura Y, Iwata I, Masuda T, Hirano Y. The neural circuit linking mushroom body parallel circuits induces memory consolidation in *Drosophila*. Proc Natl Acad Sci USA. 2019;116(32):16080–16085. doi:10.1073/pnas.1901292116

12. Bervoets S, Jacob MS, Devineni AV, et al. dArc1 controls sugar reward valuation in Drosophila melanogaster. Current Biology. 2025;35(17):4188–4198.e7. doi:10.1016/j.cub.2025.07.048

13. Lee PH, Anaya M, Ladinsky MS, Reitsma JM, Zinn K. An extracellular vesicle targeting ligand that binds to Arc proteins and facilitates Arc transport in vivo. eLife. 2023;12:e82874. doi:10.7554/eLife.82874

14. Ly S, Pack AI, and Naidoo N. The neurobiological basis of sleep: insights from Drosophila. Neurosci. Biobehav. Rev. 2018;87,67–86. 10.1016/j.neubiorev.2018.01.015.

15. Yuan Q, Joiner WJ, and Seghal A. A sleep-promoting role for the Drosophila serotonin receptor 1A. Curr. Biol. 2006;16(11)1051-1062. 10.1016/j.cub.2006.04.032.

16. Pooryasin A and Fiala A. Identified serotonin-releasing neurons induce behavioral quiescence and suppress mating in Drosophila. J. Neurosci. 2015;35(37)12792-12812. 10.1523/jneurosci.1638-15.2015.

17. Qian Y. et al. Sleep homeostasis regulated by 5HT2b receptor in a small subset of neurons in the dorsal fan-shaped body of drosophila. eLife. 2017;6,e26519. 10.7554/elife.26519.

18. Knapp EM, et al. Mutation of the Drosophila melanogaster serotonin transporter dSERT impacts sleep, courtship, and feeding behaviors. PLoS Genet. 2022;18(11):e1010289. 10.1371/journal.pgen.1010289.

19. Sitaraman D, Aso Y, Rubin GM, Nitabach MN. Control of sleep by dopaminergic inputs to the Drosophila mushroom body. Front Neural Circuits. 2015;9. doi:10.3389/fncir.2015.00073

20. Driscoll M, Buchert SN, Coleman V, McLaughlin M, Nguyen A, and Sitaraman D. Compartment specific regulation of sleep by mushroom body requires GABA and dopaminergic signaling. Sci. Rep. 2021;11(1):20067. 10.1038/s41598-021-99531-2.

21. Yan L, et al. Brief disruption of activity in a subset of dopaminergic neurons during consolidation impairs long-term memory by fragmenting sleep. bioRxiv. 2024. 10.1101/2023.10.23.563499.

22. Shafer OT, Keene AC. The regulation of Drosophila sleep. Current Biology. 2021;31(1):R38–R49. doi:10.1016/j.cub.2020.10.082

23. Shaw PJ, Cirelli C, Greenspan RJ, Tononi G. Correlates of sleep and waking in *Drosophila melanogaster*. Science. 2000;287(5459):1834–1837. doi:10.1126/science.287.5459.1834

24. Vecsey CG, Koochagian C, Porter MT, Roman G, Sitaraman D. Analysis of sleep and circadian rhythms from *Drosophila* activity-monitoring data using SCAMP. Cold Spring Harb Protoc. 2024;2024(11):pdb.prot108182. doi:10.1101/pdb.prot108182

25. Wiggin TD, Goodwin PR, Donelson NC, et al. Covert sleep-related biological processes are revealed by probabilistic analysis in *Drosophila*. Proc Natl Acad Sci USA. 2020;117(18):10024–10034. doi:10.1073/pnas.1917573117

26. Titos I, Juginović A, Vaccaro A, et al. A gut-secreted peptide suppresses arousability from sleep. Cell. 2023;186(7):1382–1397.e21. doi:10.1016/j.cell.2023.02.022

27. Chowdhury B, Abhilash L, Ortega A, Liu S, Shafer O. Homeostatic control of deep sleep and molecular correlates of sleep pressure in Drosophila. eLife. 2023;12:e91355. doi:10.7554/eLife.91355

28. Tuthill JC, Wilson RI. Mechanosensation and adaptive motor control in insects. Current Biology. 2016;26(20):R1022–R1038. doi:10.1016/j.cub.2016.06.070

29. Öztürk-Çolak A, Inami S, Buchler JR, et al. Sleep Induction by Mechanosensory Stimulation in Drosophila. Cell Reports. 2020;33(9):108462. doi:10.1016/j.celrep.2020.108462

30. Sheeba V, Fogle KJ, Kaneko M, et al. Large ventral lateral neurons modulate arousal and sleep in Drosophila. Current Biology. 2008;18(20):1537–1545. doi:10.1016/j.cub.2008.08.033

31. Au DD, Liu JC, Park SJ, et al. Drosophila photoreceptor systems converge in arousal neurons and confer light responsive robustness. Front Neurosci. 2023;17:1160353. doi:10.3389/fnins.2023.1160353

32. De Queiroz BR, Laghrissi H, Rajeev S, et al. Axonal RNA localization is essential for long-term memory. Nat Commun. 2025;16(1):2560. doi:10.1038/s41467-025-57651-7

33. Davie K, Janssens J, Koldere D, et al. A single-cell transcriptome atlas of the aging Drosophila brain. Cell. 2018;174(4):982–998.e20. doi:10.1016/j.cell.2018.05.057

34. Dopp J, Ortega A, Davie K, Poovathingal S, Baz ES, Liu S. Single-cell transcriptomics reveals that glial cells integrate homeostatic and circadian processes to drive sleep–wake cycles. Nat Neurosci. 2024;27(2):359–372. doi:10.1038/s41593-023-01549-4

35. Lee HK (Peter), Cording A, Vielmetter J, Zinn K. Interactions between a receptor tyrosine phosphatase and a cell surface ligand regulate axon guidance and glial-neuronal communication. Neuron. 2013;78(5):813–826. doi:10.1016/j.neuron.2013.04.001

36. Suzuki A, Yanagisawa M, and Greene RW. Loss of *Arc* attenuates the behavioral and molecular responses for sleep homeostasis in mice. Proc. Natl. Acad. Sci. USA. 2020;117(19):10547–10553. 10.1073/pnas.1906840117.

37. Gratz SJ, Cummings AM, Nguyen JN, et al. Genome engineering of *Drosophila* with the CRISPR RNA-guided Cas9 nuclease. Genetics. 2013;194(4):1029–1035. doi:10.1534/genetics.113.152710

38. Gratz SJ, Ukken FP, Rubinstein CD, et al. Highly specific and efficient CRISPR/Cas9-catalyzed homology-directed repair in *Drosophila*. Genetics. 2014;196(4):961–971. doi:10.1534/genetics.113.160713

39. Bervoets S, Wei N, Erfurth ML, et al. Transcriptional dysregulation by a nucleus-localized aminoacyl-tRNA synthetase associated with Charcot-Marie-Tooth neuropathy. Nat Commun. 2019;10(1):5045. doi:10.1038/s41467-019-12909-9

